# Comparing scientific abstracts generated by ChatGPT to original abstracts using an artificial intelligence output detector, plagiarism detector, and blinded human reviewers

**DOI:** 10.1101/2022.12.23.521610

**Authors:** Catherine A. Gao, Frederick M. Howard, Nikolay S. Markov, Emma C. Dyer, Siddhi Ramesh, Yuan Luo, Alexander T. Pearson

## Abstract

**Background:** Large language models such as ChatGPT can produce increasingly realistic text, with unknown information on the accuracy and integrity of using these models in scientific writing.

**Methods:** We gathered ten research abstracts from five high impact factor medical journals (n=50) and asked ChatGPT to generate research abstracts based on their titles and journals. We evaluated the abstracts using an artificial intelligence (AI) output detector, plagiarism detector, and had blinded human reviewers try to distinguish whether abstracts were original or generated.

**Results:** All ChatGPT-generated abstracts were written clearly but only 8% correctly followed the specific journal’s formatting requirements. Most generated abstracts were detected using the AI output detector, with scores (higher meaning more likely to be generated) of median [interquartile range] of 99.98% [12.73, 99.98] compared with very low probability of AI-generated output in the original abstracts of 0.02% [0.02, 0.09]. The AUROC of the AI output detector was 0.94. Generated abstracts scored very high on originality using the plagiarism detector (100% [100, 100] originality). Generated abstracts had a similar patient cohort size as original abstracts, though the exact numbers were fabricated. When given a mixture of original and general abstracts, blinded human reviewers correctly identified 68% of generated abstracts as being generated by ChatGPT, but incorrectly identified 14% of original abstracts as being generated. Reviewers indicated that it was surprisingly difficult to differentiate between the two, but that the generated abstracts were vaguer and had a formulaic feel to the writing.

**Conclusion:** ChatGPT writes believable scientific abstracts, though with completely generated data. These are original without any plagiarism detected but are often identifiable using an AI output detector and skeptical human reviewers. Abstract evaluation for journals and medical conferences must adapt policy and practice to maintain rigorous scientific standards; we suggest inclusion of AI output detectors in the editorial process and clear disclosure if these technologies are used. The boundaries of ethical and acceptable use of large language models to help scientific writing remain to be determined.

## Background

The release of OpenAI’s free tool ChatGPT^1^ on November 30, 2022 demonstrated the ability of artificial intelligence models to generate content, with articles quickly published on its possible uses and potential controversies.^2–4^ Early adopters have shared their experiences on social media, with largely positive sentiments.^5^ Articles are bemoaning the death of the traditional school essay assignment,^4,6,7^ as ChatGPT has been shown to generate high-scoring papers^8^ and even articulate critical thinking.^9^ The ethical and acceptable boundaries of ChatGPT’s use in scientific writing remain unclear.^10^

Large language models (LLM) are often complex neural network-based models that can generate tone and content-defined text. These are trained on enormous amounts of data to predict the best next text element, which generates a product that reads naturally. ChatGPT is built on Generative Pre-trained Transformer-3 (GPT-3), which is one of the largest of these types of models, trained with 175 billion parameters.^11^ These models generate coherent and fluent output, that can be difficult to distinguish from text written by humans.^12^

Artificial intelligence (AI) has numerous applications in medical technologies,^13^ and the writing of medical research is no exception, with products such as the SciNote Manuscript Writer that help with writing manuscripts.^14^ However, with the release of ChatGPT, this powerful technology is now available to all users for free, and millions are engaging with the new technology. The user base is likely to continue to grow. Thus, there is an urgent need to determine if ChatGPT can write convincing medical research abstracts.

## Methods

In this study, we evaluated the abstracts generated by ChatGPT (Version Dec 15) for 50 scientific medical papers. We gathered titles and original abstracts from current and recent issues (published in late November and December of 2022) of five high-impact journals (*JAMA, NEJM, BMJ, Lancet*, and *Nature Medicine*) and compared them with the original abstracts. The prompt fed to the model was ‘Please write a scientific abstract for the article [title] in the style of [journal] at [link]’. Note that the link is superfluous because ChatGPT cannot browse the internet. ChatGPT’s knowledge cutoff date is September 2021. Given ChatGPT is sensitive to previous queries in the same chat, we ran each prompt in a new session.

We evaluated the ChatGPT-generated abstracts for plagiarism detection using a web-crawling plagiarism detection tool,^15^ which gives an originality score of 0-100%, with 100% meaning that no plagiarism was detected. We also evaluated abstracts with an AI output detector using the GPT-2 Output Detector,^16,17^ a RoBERTa based sequence classifier which gives abstracts a score ranging from 0.02 to 99.98% ‘fake’, with a higher score indicating the text was more likely to be generated by an AI algorithm.

We evaluated whether blinded human reviewers (FMH, NSM, ECD, SR) could identify ChatGPT-generated abstracts. For every pair of reviewers, we used randomization via an electronic coin flip to decide whether an original or generated abstract would be provided for the first reviewer, with the opposite being given to the second reviewer. Each reviewer was given 25 abstracts to review, and informed that there was a mixture of original and generated abstracts in the collection. Reviewers were asked to give a binary score of whether they thought the abstract was original or generated and invited to make free-text observations while reviewing. Reviewers were not shown any data analysis until after their scoring of abstracts was completed.

We evaluated whether the format of the generated abstract adhered to the journal’s requirements by comparing it to the original article’s headings and structure. We also compared the reported patient cohort sizes between the original and generated abstracts.

Graphics and statistics were done in Python version 3.9 with *seaborn* version 0.11.2,^18^ *matplotlib* version 3.5.1,^19^ *sklearn* version 1.0.2,^20^ *scipy* version 1.7.3,^21^ and *statsannotations* version 0.4.4.^22^ Group statistics are reported using median [interquartile range] and were compared using two-sided Mann Whitney Wilcoxon (MWW) tests, with p<0.05 being the cutoff for statistical significance. Proportions were compared with Fisher’s Exact tests. Correlation between the cohort sizes was done with Pearson’s correlation.

## Results

We asked ChatGPT to generate 50 scientific abstracts in the format of specific journals using the prompt above (example subset in Supplemental Materials). While all the output appeared superficially to be formatted as a scientific abstract, only 8 (16%) correctly used the headings particular to the specific journal in the prompt (e.g., *Nature Medicine’s* paragraph-style without headings, as opposed to specific headings such as ‘Design, Setting, and Participants’ for *JAMA*, see Appendix for examples). The patient cohort sizes were a similar order of magnitude between the original abstracts and the generated abstracts, with a Pearson correlation of the logarithmic cohort sizes of r=0.76, p<0.001 (Figure 1).

**Figure 1.**
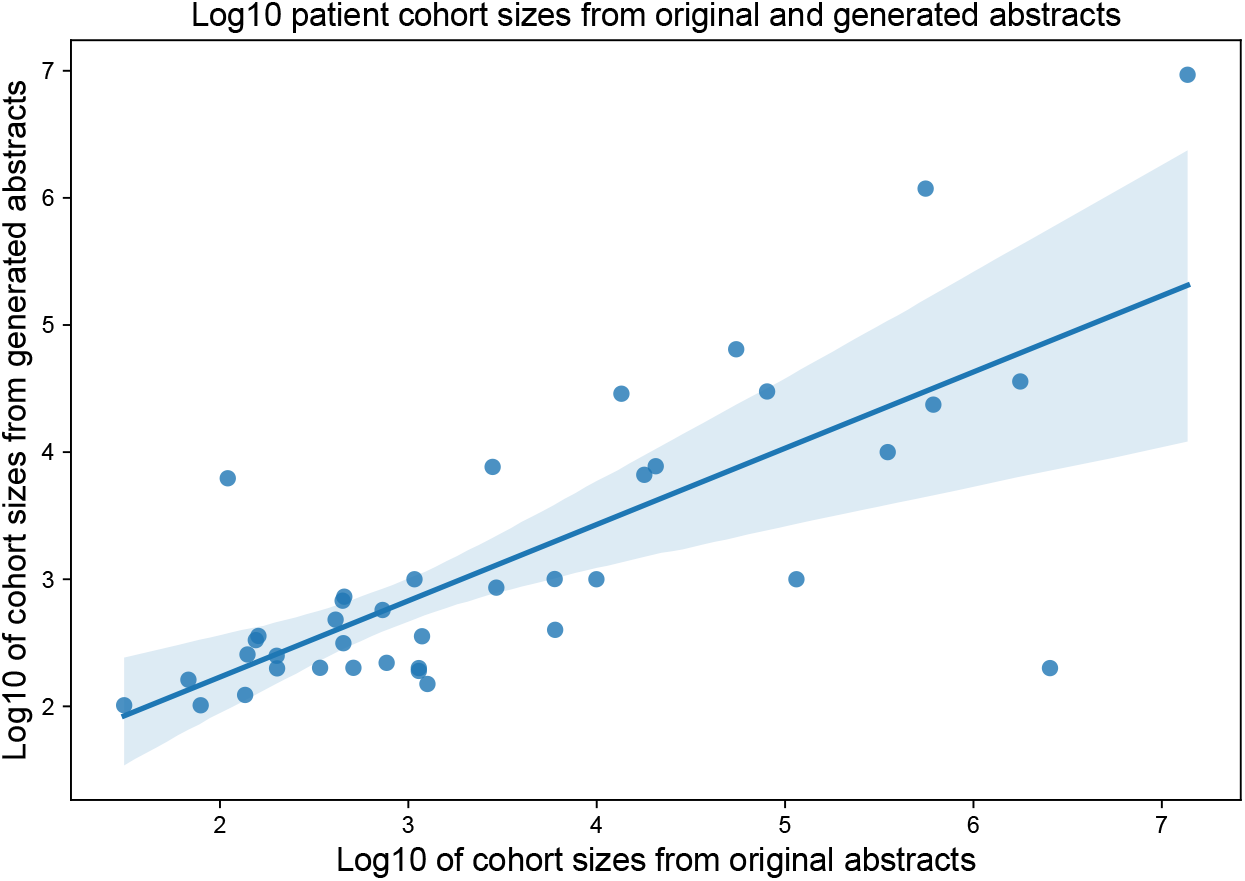
Generated abstracts have a similar patient cohort size as original abstracts. Cohort sizes from original abstracts (x-axis) and generated abstracts (y-axis) plotted on a logarithmic 10 scale.

The AI output detector found a high probability (higher AI detection score % ‘fake’ being more likely to be generated text) of AI-generated output in many of the generated abstracts median [IQR] of 99.98% [12.73,99.98] compared with very low probability of AI-generated output in nearly all the original abstracts 0.02% [0.02,0.09] (Figure 2a). However, 17 (34%) generated abstracts received a score below 50% from the AI output detector, including 5 with scores below 1%. There was only one (2%) original abstract that scored above 50% on the AI output detector. The AI output detector had an area under the receiver operating characteristics (AUROC) curve of 0.94, indicating excellent discrimination between original and generated abstracts (Figure 2b).

**Figure 2.**
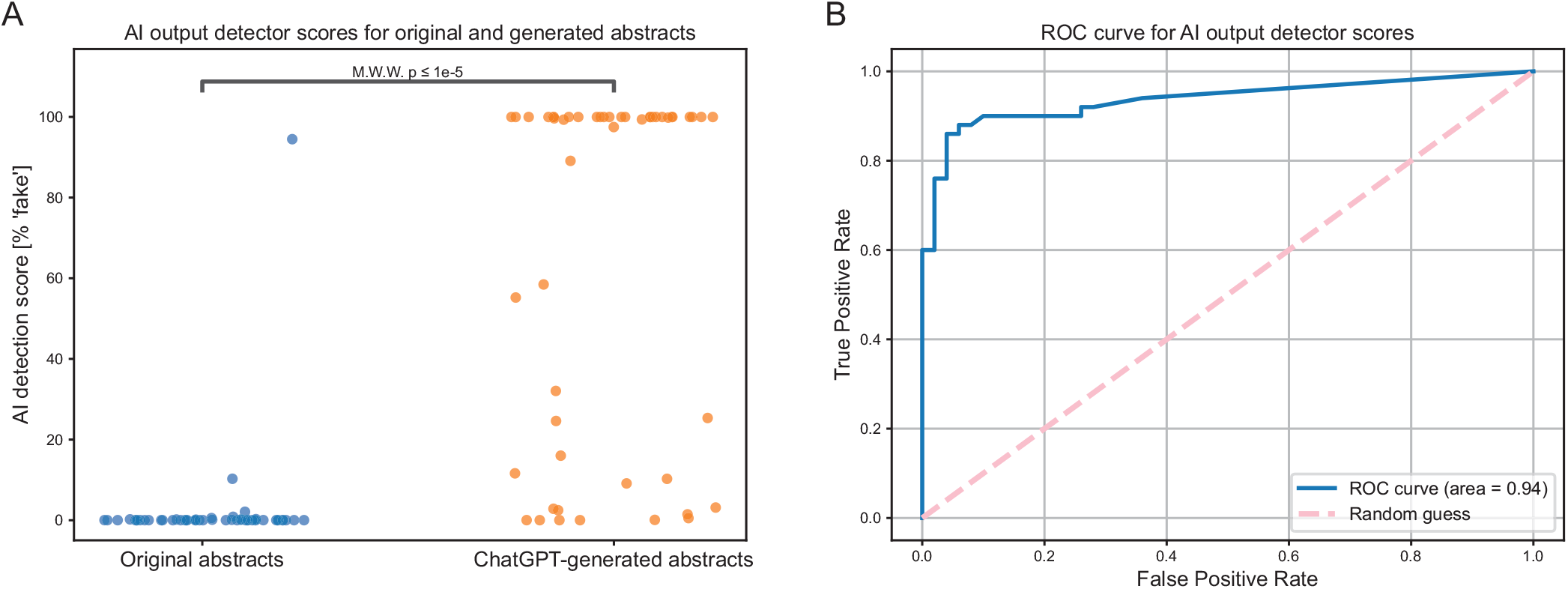
Many generated abstracts can be detected using an AI output detector. **(A)** AI detection scores as [% ‘fake’] per GPT-2 Output Detector for original abstracts and generated abstracts. Except for one case, all original abstracts scored extremely low on the AI output detector. The majority of generated abstracts scored high on the AI output detector, but 17 (34%) scored lower than 50%. **(B)** The AI output detector ROC curve for discriminating between original and generated abstracts, with AUROC of 0.94.

Nearly all the generated text were deemed completely original by the plagiarism checker with a median originality score of 100% [IQR 100,100] (Figure 3). As a positive control, we ran original articles through the plagiarism checker, with originality scores of 38.5% [15.25,56.75], almost always matching against the original article as the source of ‘plagiarism’.

**Figure 3.**
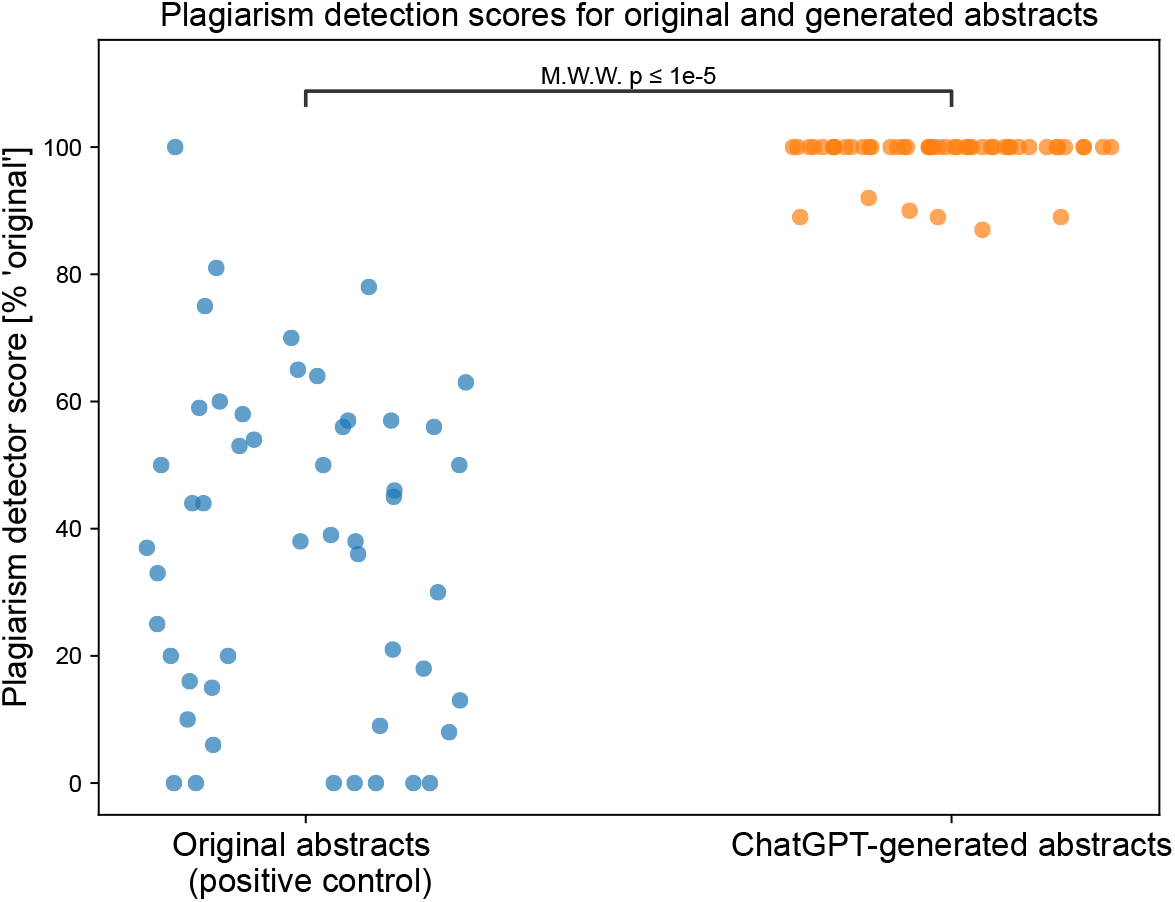
Generated abstracts are original and do not plagiarize from other written work. Plagiarism checker report for original abstracts and generated abstracts [% original] (higher value meaning fewer matching text was found).

Reviewers were able to correctly identify 68% of generated abstracts as being generated, and correctly identified 86% of original articles as being original. They incorrectly identified 32% of generated abstracts as being real, and 14% of original abstracts as being generated (p<0.001 by Fisher’s Exact test).

**Table 1.** Reviewers were correct in differentiating original versus generated abstracts most of the time. Human reviewer scoring for whether abstracts were real or generated, along with truth.

|  |  | Truth |  |
| --- | --- | --- | --- |
|  |  | Original | Generated |
| Reviewer guess | Original | 43 | 16 |
|  | Generated | 7 | 34 |

Our reviewers commented that abstracts they thought were generated by ChatGPT were superficial and vague, and focused on details of original abstracts such as inclusion of Clinical Trial Registration numbers and alternative spellings of words. Observations noted while interacting with the model were its use of generated numbers, including (nonexistent) clinical trial number.

The AI detection scores were not statistically different (p=0.45 by MWW) between the abstracts that reviewers correctly identified as generated and ones that they failed to identify as generated (Figure 4).

**Figure 4.**
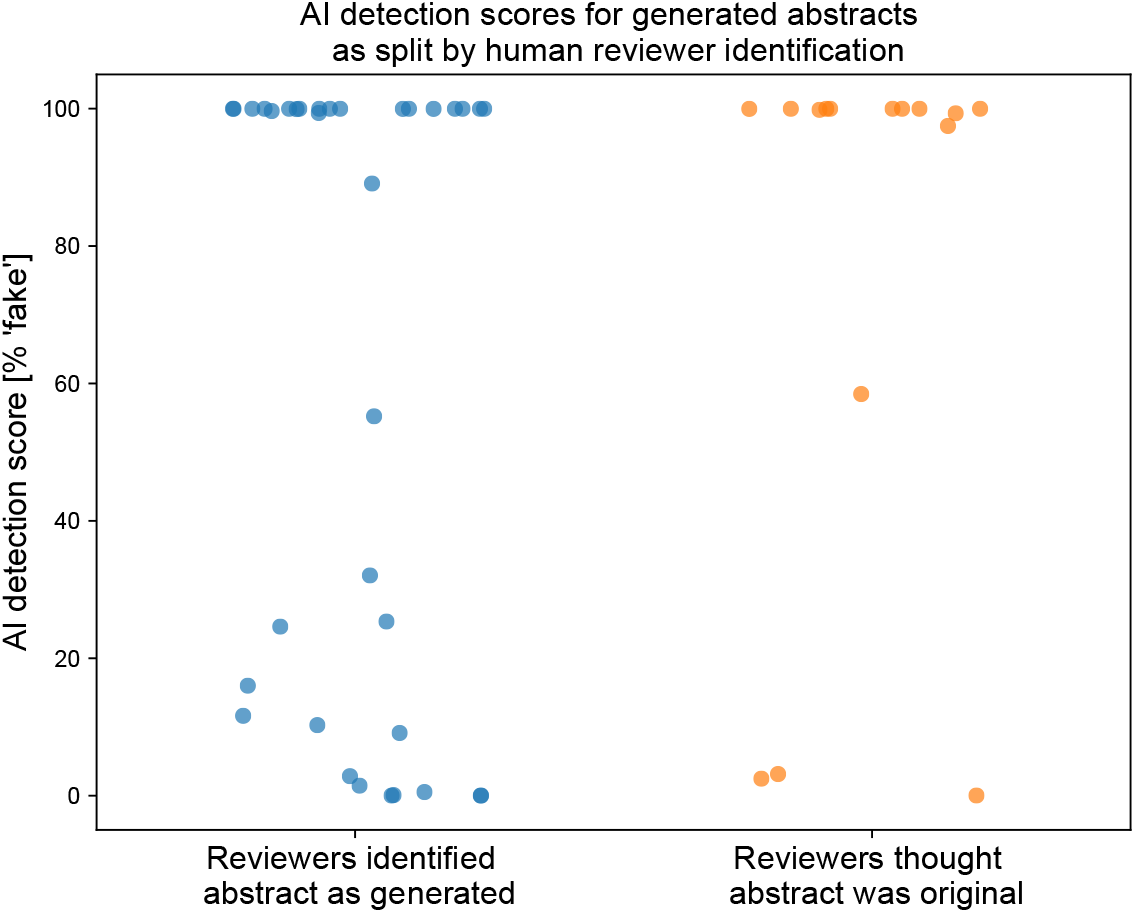
Reviewers use criteria different than the AI output detector for flagging abstracts as either generated or original. The AI detection scores for generated abstracts were not different between abstracts that human reviewers identified as generated, and those that they failed to identify as generated.

## Discussion

In this study, we found that both humans and AI-output detectors were able to identify abstracts generated by ChatGPT in the majority of cases, but neither were perfect discriminators. Our reviewers misclassified a portion of real abstracts as being generated, indicating they were highly skeptical when reviewing the abstracts. It was impressive that given only a title and journal, ChatGPT was able to generate a superficially readable abstract, with accurate themes and topic-specific patient cohort sizes. However, the actual numbers in the abstract were fabricated, and the model was only able to correctly format the abstract to journal specifications a minority of the time. This is the first study to our knowledge to evaluate the ability of the new ChatGPT model to write convincing medical research abstracts using AI output detectors, plagiarism detectors, and blinded human reviewers.

Limitations to our study include its small sample size and few reviewers. ChatGPT is also known to be sensitive to small changes in prompts; we did not exhaust different prompt options, nor did we deviate from our prescribed prompt. ChatGPT generates a different response even to the same prompt multiple times, and we only evaluated one of infinite possible outputs. The maximum input for the AI output detector is 510 tokens, thus some of the abstracts were not able to be fully evaluated due to their length. Our human reviewers knew that a subset of the abstracts they were viewing were generated by ChatGPT, but a reviewer outside this context may not be able to distinguish them as written by a large language model. Future studies could expand on our sample size and methodology to include other AI output detector models.

We anticipate that this technology could be used in both an ethical and unethical way. Given its ability to generate abstracts with believable numbers, it could be used to entirely falsify research. On the other hand, the technology may be used in conjunction with a researcher’s own scientific knowledge as a tool to decrease the burden of writing and formatting. It could be used by scientists publishing in a language that is not their native language, to improve equity. Furthermore, AI models have been shown to be highly sensitive to biases in training data,^23,24^ and further data is needed to determine the potential for bias perpetuated by ChatGPT - especially given the overt prejudices emerging from prior language generation models.^25^

We suggest clear disclosure when a manuscript is written with assistance from ChatGPT;^26^ some have even included it as a co-author.^27^ Reassuringly, there are patterns that allow it to be detected by AI output detectors. Though there is ongoing work to embed watermarks in output, until this is standardized and robust against scrubbing, we suggest running journal and conference abstract submissions through AI output detectors as part of the research editorial process to protect from targeting by organizations such as paper mills.

Abstract generation by ChatGPT is a powerful tool to create readable scientific abstracts. The generated abstracts do not alarm plagiarism-detection models, as the text is generated anew, but can often be detected using AI detection models, and identified by a blinded human reviewer. The optimal use and ethical boundaries of AI-generated writing remain to be determined.

## Data availability

The data used in the manuscript are available upon reasonable request to the corresponding author.

## Contributions

CAG and FMH conceived and designed the study, with contributions from NSM, YL, ATP.

CAG acquired, generated, and analyzed the data.

FMH, NSM, ECD, SR reviewed the abstracts.

CAG drafted the manuscript. All authors reviewed the data and revised the manuscript.

## Supplemental Materials

An example subset of the generated abstracts.

“Association of COVID-19 Vaccinations With Intensive Care Unit Admissions and Outcome of Critically Ill Patients With COVID-19 Pneumonia in Lombardy, Italy” in the style of JAMA; original paper^28^

### Background

COVID-19 has had a significant impact on healthcare systems worldwide, leading to high rates of intensive care unit (ICU) admissions and mortality. Vaccines have been developed and deployed as a means to combat the pandemic, but their effect on critically ill patients with COVID-19 pneumonia remains unclear.

### Objective

To determine the association of COVID-19 vaccinations with ICU admissions and outcomes in critically ill patients with COVID-19 pneumonia in Lombardy, Italy.

### Design

Retrospective cohort study.

### Setting

ICUs in Lombardy, Italy.

### Participants

A total of 314 critically ill patients with COVID-19 pneumonia admitted to ICUs in Lombardy between January 1 and June 30, 2021.

### Exposure

COVID-19 vaccination status.

### Main Outcomes and Measures

The primary outcome was ICU admission, and secondary outcomes included ICU mortality and length of ICU stay.

### Results

Of the 314 critically ill patients with COVID-19 pneumonia, 149 (47.5%) were vaccinated. The vaccinated group had a significantly lower rate of ICU admission compared with the unvaccinated group (47.0% vs 63.2%; difference, −16.2% [95% CI, −26.6% to −5.9%]; P = .002). In addition, vaccinated patients had a lower ICU mortality rate compared with unvaccinated patients (14.8% vs 25.0%; difference, −10.2% [95% CI, −20.8% to 0.3%]; P = .05). The length of ICU stay did not differ significantly between the vaccinated and unvaccinated groups (median, 8 days [interquartile range, 5-13 days] vs 9 days [interquartile range, 5-14 days]; P = .50).

### Conclusions and Relevance

In this cohort of critically ill patients with COVID-19 pneumonia, vaccination was significantly associated with lower rates of ICU admission and mortality. These findings suggest that COVID-19 vaccines may provide a potentially important benefit for critically ill patients with COVID-19 pneumonia.

“Once-Weekly Semaglutide in Adolescents with Obesity” in the style of NEJM; original paper^29^

### Objective

To evaluate the efficacy and safety of once-weekly semaglutide, a glucagon-like peptide-1 receptor agonist, in adolescents with obesity.

### Methods

This was a randomized, double-blind, placebo-controlled, parallel-group trial conducted at 29 clinical centers in the United States and Canada. A total of 199 adolescents aged 12 to <18 years with a body mass index ≥35 kg/m2 or ≥30 kg/m2 with comorbidities were randomized to receive once-weekly subcutaneous injections of semaglutide (1.0 or 0.5 mg) or placebo for 68 weeks. The primary outcome was the change in body weight from baseline to week 68.

### Results

At week 68, the mean change in body weight was −11.9 kg in the semaglutide 1.0-mg group, −9.5 kg in the semaglutide 0.5-mg group, and −2.5 kg in the placebo group (P < 0.001 for both semaglutide groups vs. placebo). The proportion of participants who lost ≥5% and ≥10% of their body weight was greater in the semaglutide groups than in the placebo group (P < 0.001 for both comparisons). Semaglutide was associated with statistically significant improvements in blood pressure, glycemic control, and lipid profiles. The incidence of adverse events was similar in the semaglutide and placebo groups, with no significant differences in the incidence of serious adverse events or serious treatment-emergent adverse events.

### Conclusions

In this randomized trial, once-weekly semaglutide was effective and well tolerated in adolescents with obesity, leading to clinically meaningful and statistically significant weight loss and improvements in cardiometabolic risk factors.

### Trial Registration

ClinicalTrials.gov, NCT03702497.

“Chlorthalidone vs. Hydrochlorothiazide for Hypertension–Cardiovascular Events” in the style of NEJM; original article^30^

### Background

Thiazide diuretics are commonly used as first-line agents for the management of hypertension. Chlorthalidone and hydrochlorothiazide are the two most commonly prescribed thiazides in the United States, but their comparative effectiveness and safety in reducing cardiovascular events has not been fully established.

### Methods

We conducted a multicenter, randomized, double-blind, controlled trial to compare the effects of chlorthalidone and hydrochlorothiazide on blood pressure control and cardiovascular outcomes in adults with hypertension. Eligible participants were randomly assigned to receive either chlorthalidone (12.5 to 25 mg/day) or hydrochlorothiazide (12.5 to 50 mg/day) for a median follow-up of 4.9 years. The primary outcome was a composite of major cardiovascular events (myocardial infarction, stroke, or death from cardiovascular causes).

### Results

A total of 28,803 participants were enrolled, 14,413 in the chlorthalidone group and 14,390 in the hydrochlorothiazide group. The mean age was 61.9 years, and 63.7% were women. The mean baseline systolic blood pressure was 152.7 mm Hg. The two groups had similar blood pressure control and demographic characteristics. The incidence of the primary outcome was significantly lower in the chlorthalidone group than in the hydrochlorothiazide group (9.1% vs. 11.1%; hazard ratio, 0.81; 95% confidence interval [CI], 0.74 to 0.88; P<0.001). The incidence of myocardial infarction was also significantly lower in the chlorthalidone group than in the hydrochlorothiazide group (4.4% vs. 5.7%; hazard ratio, 0.78; 95% CI, 0.68 to 0.88; P<0.001). There were no significant between-group differences in the incidence of stroke or death from cardiovascular causes. Adverse events were similar in the two groups.

### Conclusions

In this large, randomized, controlled trial, chlorthalidone was superior to hydrochlorothiazide in reducing the incidence of major cardiovascular events in adults with hypertension. These findings support the use of chlorthalidone as a preferred treatment option for hypertension.

“Racial disparities in recommendations for surgical resection of primary brain tumours: a registry-based cohort analysis” in the style of The Lancet

### Introduction

Despite progress in the treatment of primary brain tumours, racial disparities in patient outcomes persist. Previous studies have suggested that these disparities may be related to differences in treatment recommendations and access to care. However, the extent to which racial disparities exist in the recommendation of surgical resection for primary brain tumours is unclear.

### Methods

We conducted a registry-based cohort analysis using data from the National Cancer Database to examine racial disparities in recommendations for surgical resection of primary brain tumours in the United States. We included patients diagnosed with a primary brain tumour between 2010 and 2014, and excluded those with metastatic disease or missing race/ethnicity data. We compared the proportion of white and nonwhite patients who received a recommendation for surgical resection, adjusting for patient, tumour, and hospital characteristics.

### Results

Among the 23,632 patients included in the analysis, 63.4% were white and 36.6% were non-white. After adjusting for patient, tumour, and hospital characteristics, non-white patients were significantly less likely to receive a recommendation for surgical resection than white patients (odds ratio 0.87, 95% CI 0.81-0.93).

### Conclusion

Our study found significant racial disparities in the recommendation of surgical resection for primary brain tumours in the United States. Further research is needed to identify the causes of these disparities and to develop interventions to reduce them.

“Efficacy of awake prone positioning in patients with covid-19 related hypoxemic respiratory failure: systematic review and meta-analysis of randomized trials” in the style of The BMJ, original article^31^

### Objective

To evaluate the efficacy of awake prone positioning (APP) in patients with COVID-19 related hypoxemic respiratory failure.

### Design

Systematic review and meta-analysis of randomized controlled trials (RCTs).

### Data sources

A comprehensive search was conducted in multiple databases including PubMed, Embase, and the Cochrane Library, up to June 2021.

### Eligibility criteria

RCTs that compared APP with standard care in patients with COVID-19 related hypoxemic respiratory failure were included.

### Data extraction and synthesis

Two reviewers independently extracted data and assessed the risk of bias. The primary outcome was oxygenation index (OI), defined as the ratio of partial pressure of arterial oxygen (PaO2) to fraction of inspired oxygen (FiO2). The secondary outcomes were 28-day mortality, duration of mechanical ventilation, and length of hospital stay. The meta-analysis was performed using a fixed-effect model.

### Results

Six RCTs with a total of 859 patients were included in the final analysis. The meta-analysis showed that APP significantly improved OI compared to standard care (mean difference [MD] −10.65, 95% confidence interval [CI] −15.29 to −5.99, p < 0.001). There was also a trend towards a lower 28-day mortality in the APP group (relative risk [RR] 0.63, 95% CI 0.40 to 0.98, p = 0.04). There were no significant differences in the duration of mechanical ventilation (MD −1.40, 95% CI −3.90 to 1.11, p = 0.27) or length of hospital stay (MD - 2.43, 95% CI −6.69 to 1.82, p = 0.28) between the two groups.

### Conclusion

This systematic review and meta-analysis suggests that APP may be effective in improving oxygenation and reducing 28-day mortality in patients with COVID-19 related hypoxemic respiratory failure. Further high-quality RCTs are needed to confirm these findings.

“Atezolizumab plus anthracycline-based chemotherapy in metastatic triple-negative breast cancer: the randomized, double-blind phase 2b ALICE trial” in the style of Nature Medicine; original article^32^

Triple-negative breast cancer (TNBC) is a subtype of breast cancer that is aggressive and difficult to treat. Atezolizumab is a monoclonal antibody that targets the protein PD-L1, which is expressed on some cancer cells and can help them evade the immune system. In this double-blind, phase 2b clinical trial, called ALICE, we evaluated the safety and efficacy of adding atezolizumab to an anthracycline-based chemotherapy regimen in patients with metastatic TNBC. A total of 162 patients were randomized to receive either atezolizumab plus chemotherapy or placebo plus chemotherapy. The primary endpoint was progression-free survival (PFS), which was significantly longer in the atezolizumab group compared to the placebo group (median PFS of 6.3 months vs 3.7 months, respectively; hazard ratio 0.59, 95% confidence interval 0.43-0.81, p=0.0007). Overall survival and objective response rate were also higher in the atezolizumab group, although the differences did not reach statistical significance. Adverse events were similar between the two groups, with the most common being neutropenia, anemia, and nausea. In conclusion, the addition of atezolizumab to anthracycline-based chemotherapy significantly improves PFS in patients with metastatic TNBC, with a manageable safety profile. These findings support the further investigation of atezolizumab in this patient population.

## Appendix

Formatting header specifications noted for journals JAMA, NEJM, BMJ, Lancet, and Nature Medicine.

### JAMA

Importance

Objective

Design, Setting, and Participants

Interventions

Main Outcomes and Measures

Results

Conclusions and Relevance

### The NEJM

Background

Methods

Results

Conclusions

### The BMJ (specific headers variable depending on type of study)

Objective

Design

Data source

Setting

Participants

Data extraction and synthesis

Main outcome measures

Results

Conclusions

### The Lancet

Background

Methods

Findings

Interpretation

*Nature Medicine* - headerless paragraph abstract

## Works Cited

1. OpenAI. ChatGPT: Optimizing language models for dialogue. OpenAI. Published November 30, 2022. Accessed December 17, 2022. https://openai.com/blog/chatgpt/

2. Shankland S. ChatGPT: Why everyone is obsessed this mind-blowing AI chatbot. CNET. Published December 14, 2022. Accessed December 18, 2022. https://www.cnet.com/tech/computing/chatgpt-why-everyone-is-obsessed-this-mind-blowing-ai-chatbot/

3. Agomuoh F. ChatGPT: how to use the viral AI chatbot that took the world by storm. Digital Trends. Published December 13, 2022. Accessed December 18, 2022. https://www.digitaltrends.com/computing/how-to-use-openai-chatgpt-text-generation-chatbot/

4. Hern A. AI bot ChatGPT stuns academics with essay-writing skills and usability. The guardian. https://www.theguardian.com/technology/2022/dec/04/ai-bot-chatgpt-stuns-academics-with-essay-writing-skills-and-usability. Published December 4, 2022. Accessed December 18, 2022.

5. Haque MU, Dharmadasa I, Sworna ZT, Rajapakse RN, Ahmad H. “I think this is the most disruptive technology”: Exploring Sentiments of ChatGPT Early Adopters using Twitter Data. arXiv [csCL]. Published online December 12, 2022. http://arxiv.org/abs/2212.05856

6. Stokel-Walker C. AI bot ChatGPT writes smart essays - should professors worry? Nature. Published online December 9, 2022. doi:10.1038/d41586-022-04397-7

7. Whitford E. Here’s how Forbes got the ChatGPT AI to write 2 college essays in 20 minutes. Forbes. Published December 9, 2022. Accessed December 18, 2022. https://www.forbes.com/sites/emmawhitford/2022/12/09/heres-how-forbes-got-the-chatgpt-ai-to-write-2-college-essays-in-20-minutes/?sh=2b5a552456ad

8. Yeadon W, Inyang OO, Mizouri A, Peach A, Testrow C. The death of the short-form Physics essay in the coming AI revolution. arXiv [physics.ed-ph]. Published online December 22, 2022. doi:10.48550/ARXIV.2212.11661

9. Susnjak T. ChatGPT: The end of online exam integrity? arXiv [csAI]. Published online December 19, 2022. http://arxiv.org/abs/2212.09292

10. Much to discuss in AI ethics. Nat Mach Intell. 2022;4(12):1055–1056.

11. Korngiebel DM, Mooney SD. Considering the possibilities and pitfalls of Generative Pre-trained Transformer 3 (GPT-3) in healthcare delivery. NPJ Digit Med. 2021;4(1):93.

12. Clark E, August T, Serrano S, Haduong N, Gururangan S, Smith NA. All that’s ‘human’ is not gold: Evaluating human evaluation of generated text. In: Proceedings of the 59th Annual Meeting of the Association for Computational Linguistics and the 11th International Joint Conference on Natural Language Processing (Volume 1: Long Papers). Association for Computational Linguistics; 2021:7282–7296.

13. Briganti G, Le Moine O. Artificial intelligence in medicine: Today and tomorrow. Front Med (Lausanne). 2020;7:27.

14. Masa. SciNote Manuscript Writer - using Artificial Intelligence. SciNote. Published November 18, 2020. Accessed December 17, 2022. https://www.scinote.net/blog/scinote-can-write-draft-scientific-manuscript-using-artificial-intelligence/

15. Arcade C. Plagiarism checker. Plagiarismdetector.net. Published December 28, 2010. Accessed December 17, 2022. https://plagiarismdetector.net/

16. Solaiman I, Brundage M, Clark J, et al. Release strategies and the social impacts of language models. arXiv [csCL]. Published online August 24, 2019. Accessed December 23, 2022. https://d4mucfpksywv.cloudfront.net/papers/GPT_2_Report.pdf

17. GPT-2 Output Detector. Accessed December 17, 2022. https://huggingface.co/openai-detector

18. Waskom M. seaborn: statistical data visualization. J Open Source Softw. 2021;6(60):3021.

19. Hunter JD. Matplotlib: A 2D Graphics Environment. Comput Sci Eng. 2007;9(3):90–95.

20. Pedregosa F, Varoquaux G, Gramfort A, et al. Scikit-learn: Machine Learning in Python. J Mach Learn Res. 2011;12(85):2825–2830.

21. Virtanen P, Gommers R, Oliphant TE, et al. SciPy 1.0: fundamental algorithms for scientific computing in Python. Nat Methods. 2020;17(3):261–272.

22. Charlier F, Weber M, Izak D, et al. Trevismd/Statannotations: V0.5. Zenodo; 2022. doi:10.5281/ZENODO.7213391

23. Howard FM, Dolezal J, Kochanny S, et al. The impact of site-specific digital histology signatures on deep learning model accuracy and bias. Nat Commun. 2021;12(1):4423.

24. Banerjee I, Bhimireddy AR, Burns JL, et al. Reading race: AI recognises patient’s racial identity in medical images. arXiv [csCV]. Published online July 21,2021. http://arxiv.org/abs/2107.10356

25. Bishop JM. Artificial intelligence is stupid and causal reasoning will not fix it. Front Psychol. 2020;11:513474.

26. Blanco-Gonzalez A, Cabezon A, Seco-Gonzalez A, et al. The role of AI in drug discovery: Challenges, opportunities, and strategies. arXiv [csCL]. Published online December 8, 2022. http://arxiv.org/abs/2212.08104

27. Kung TH, Cheatham M, Medenilla A, et al. Performance of ChatGPT on USMLE: Potential for AI-assisted medical education using large language models. bioRxiv. Published online December 20, 2022:2022.12.19.22283643. doi:10.1101/2022.12.19.22283643

28. Grasselli G, Zanella A, Carlesso E, et al. Association of COVID-19 vaccinations with intensive care unit admissions and outcome of critically ill patients with COVID-19 pneumonia in Lombardy, Italy. JAMA Netw Open. 2022;5(10):e2238871.

29. Weghuber D, Barrett T, Barrientos-Pérez M, et al. Once-weekly semaglutide in adolescents with obesity. N Engl J Med. 2022;387(24):2245–2257.

30. Ishani A, Cushman WC, Leatherman SM, et al. Chlorthalidone vs. Hydrochlorothiazide for hypertension-cardiovascular events. N Engl J Med. Published online December 14, 2022. doi:10.1056/NEJMoa2212270

31. Weatherald J, Parhar KKS, Al Duhailib Z, et al. Efficacy of awake prone positioning in patients with covid-19 related hypoxemic respiratory failure: systematic review and meta-analysis of randomized trials. BMJ.2022;379:e071966.

32. Røssevold AH, Andresen NK, Bjerre CA, et al. Atezolizumab plus anthracycline-based chemotherapy in metastatic triple-negative breast cancer: the randomized, double-blind phase 2b ALICE trial. Nat Med. Published online December 8, 2022:1–11.

